# First nationwide mosquito surveillance in Northern Irish wetlands reveals presence of *Culex pipiens*/*molestus* hybrids

**DOI:** 10.64898/2026.10.08.757519

**Authors:** Ryan M. Carmichael, Georgia Kirby, Luca Sloan, Aidan Desjardins, Jean-Philippe Parvy, Francesco Baldini, Emilie Pondeville, Jaimie T.A. Dick, Heather M. Ferguson, Ross N. Cuthbert

**Affiliations:** Institute for Global Food Security, School of Biological Sciences, Queen’s University Belfast, BT9 7BL, UK; MRC-University of Glasgow Centre for Virus Research, 464 Bearsden Road, G61 1QH, UK; School of Biodiversity, One Health and Veterinary Medicine, University of Glasgow, Graham Kerr Building, Glasgow, G12 8QQ, UK

**Keywords:** vector ecology, surveillance, Culicidae, bridge vector, zoonosis

## Abstract

While mosquito invasions and associated pathogens are increasingly prominent in Europe, evidence gaps remain for many northerly regions where proactive management interventions may be most effective. At the regional periphery of Europe, there remains a significant knowledge gap concerning mosquito vector distributions in Northern Ireland. This study aimed to address this major gap by systematically surveying larval and adult mosquito populations in key habitats (ponds, reedbeds, saltmarsh, wet grasslands, wet woodlands, and urban gardens). We further identified environmental drivers of important vector species abundance and screened collected specimens with molecular techniques to identify presence of cryptic human-biting vector species. In Northern Ireland, mosquitoes (including potential bridge vectors of West Nile and Usutu viruses) were found to be widely distributed. Between May and August 2025, a total of 2430 mosquitoes (951 adults and 1479 larvae) were collected over 196 trap nights and 76 larval collections. Collections encompassed 17 species, with *Culex pipiens/torrentium* being the most widely distributed and abundant taxon recorded. Mosquito abundances were greatest in reedbed habitats, related negatively with rainfall and peaked in June and July. A subset of *Culex pipiens/torrentium* mosquitoes were identified to species/biotype level using PCR, with the majority (89.3%) identified as *Culex pipiens pipien*s and 9.3% as *Culex torrentium*. Importantly, the first detections of *Culex pipiens/molestus* hybrids (1.8%) on the island of Ireland were found. This work underscores the wide, yet previously undocumented abundance and distribution of potential zoonotic vectors in Northern Ireland and provides an important baseline for management planning and future research, while contributing to broader understandings of environmental population drivers and distributions.

## Introduction

Mosquito-borne diseases (MBDs) are a growing threat in Europe due to a myriad of environmental changes including biological invasion, warming climate and anthropogenic land use (Cuthbert et al., 2023). Climate change in Europe is enhancing the transmission capacity of both native and invasive vector species, with warming temperatures favouring pathogen replication and shorter incubation times in mosquitoes (Semenza & Paz, 2021). Increasingly, native mosquito vectors such as *Culex pipiens* (Linnaeus) have played roles in the transmission of arboviruses in Europe like Usutu virus (USUV) (Chen et al., 2024) and West Nile virus (WNV) (Koch et al., 2024).

The United Kingdom of Great Britain & Northern Ireland (hereafter, UK) harbours inherent vector and pathogen biosecurity benefits as a group of islands off continental Europe. This geographic positioning reduces the risk of invasive mosquito vector establishment and disease spread, though there have been recurrent introductions of *Aedes albopictus* (Skuse) in England since 2016 (Johnston et al., 2025; Medlock et al., 2017). However, concurrent to the incursion and establishment of invasive mosquito vectors is the introduction of pathogens via infected animal hosts (*e.g.*, migratory birds), which can be transmitted by native mosquito species (Kirby et al., 2025). Indeed, migratory birds could introduce pathogens from Africa and Europe to native mosquito populations in the UK, as seen with the geographical expansion of both USUV and WNV throughout Europe associated with migratory bird routes (Engel et al., 2016; Lu et al., 2024). USUV was detected for the first time in the UK in a blackbird in London, 2020 (Folly et al., 2020), with *Cx. pipiens* mosquitoes suspected to be the main vector responsible (Lawson et al., 2022) and has expanded most recently to Scotland in 2025 (Ionescu et al., 2026). Notably, WNV was recently detected in *Ae. vexans* (Meigen) mosquitoes in England for the first time in 2023 (Bruce et al., 2025). Whilst USUV does not have high morbidity in human hosts (Cadar & Simonin, 2023), it causes severe illness in avian hosts (Agliani et al., 2025) and has been associated with declines in blackbird populations (Hanmer et al., 2026). WNV is contrastingly associated with severe neurological disease in human patients (Byas & Ebel, 2020), thus its potential emergence is of particular concern for public health.

*Culex pipiens* is a common and widespread European species complex which inhabits wetlands (*e.g.*, pools and ponds) and urban habitats (*e.g.*, artificial containers) (Medlock et al., 2005), facilitating coexistence with human habitations (Krol et al., 2024). The *Culex pipiens* complex is widely distributed throughout England, Wales (Widlake et al., 2025) and Scotland (Kirby et al., 2026). Although considered a single species, *Cx. pipiens* encompasses two morphologically identical but ecologically distinct biotypes: *Cx. pipiens pipiens* and *Cx. pipiens molestus* (Aardema et al., 2022). *Culex pipiens pipiens* is primarily ornithophilic, while *Cx. pipiens molestus* is primarily mammalophilic (Aardema et al., 2022). Hybridisation between biotypes may give rise to mosquitoes with generalist feeding preferences, but the extent of potential hybridisation is spatially variable and unevenly assessed in the UK context (Widlake et al., 2025). The morphologically-identical sister species, *Culex torrentium* (Martini), may also play a role in disease transmission (*e.g.*, (Jansen et al., 2019, 2023)), though due to morphological similarity, it is frequently grouped together with *Cx. pipiens* in arbovirus surveillance (Gobbo et al., 2025). Therefore, the role of *Cx. torrentium* in disease transmission cycles is not fully understood. In addition to *Cx. pipiens*, other native mosquito species could play a role as disease vectors in the UK and are associated with a diverse range of wetland habitats, including *Aedes detritus* (Haliday), a saltmarsh-breeding mosquito which could is a competent vector of WNV (Blagrove et al., 2016), and potential bridge vector *Culiseta annulata* (Edwards), which breeds in both wetlands and artificial habitats and feeds on both mammalian and avian hosts (Medlock et al., 2005).

Historically, mosquito vector surveillance efforts in the UK have focused on England and Wales, with little known about the contemporary distribution of mosquitoes elsewhere in theUK. Recent surveys have updated species records in Scotland, including the presence of bridge vector *Cx. p. molestus* (Kirby et al., 2026). There has remained a knowledge gap in Northern Ireland, for which the last widespread record of mosquito vector distribution was published over 30 years ago, and was itself based on historical and museum records from the 19^th^ and 20^th^ centuries (Ashe et al., 1991). Snapshot surveillance in the Republic of Ireland in 2017-18 has since updated the species record on the island, confirming the presence of three cryptic taxa: *Cx. torrentium*, *Anopheles daciae* (Linton, Nicolescu & Harbach) and *Ae. geminus* (Peus) (Schaffner et al., 2024). Furthermore, DNA barcoding has verified the occurrence of *Cx. torrentium* in Northern Ireland (Curran et al., 2025). However, no studies to-date have systematically sampled mosquitoes in Irish wetlands to assess their distribution and habitat associations, as well as environmental predictors of their presence and abundance. Further, despite presence in Great Britain (Kirby et al., 2026; Widlake et al., 2025), there has been no detection of the presence of *Cx. p. molestus*, and by extension, hybrid forms on the island of Ireland. Without an updated knowledge and understanding of mosquito vector diversity, distribution, density, and factors influencing these, Northern Ireland is left underprepared for the potential emergence of MBDs. To the best of our knowledge, the only ongoing mosquito surveillance programme in Northern Ireland focuses on sea and air ports in Belfast, and only targets *Ae. albopictus* (Vaux & Medlock, 2015).

Therefore, we conducted a comprehensive field study to fill this gap by (1) surveying larval and adult populations of mosquitoes systematically across six key habitat types during the mosquito active season in 2025, to determine species diversity, community composition and assess for the possible introduction of non-native vectors; (2) identifying potential environmental drivers of mosquito presence and abundance; and (3) testing collected specimens of *Culex* by PCR to investigate the potential presence of key bridge vector *Cx. p. molestus* and *pipiens/molestus* hybrids, and identify associated habitats.

## Methods

### Study design

Eighteen sites were systematically assessed for mosquito diversity and abundance once per month between May and August 2025, identified as containing one of six target habitat types for investigation due to their functional association with unique mosquito taxa (Medlock et al., 2024): ponds, reedbed, saltmarsh, wet grassland, wet woodland and urban gardens that contained pre-existing artificial containers filled with water (Figure 1, Table S1). Adult trapping specifically targeted these habitat types by deployment of traps in close proximity (≤ 50 metres) to the focal habitat, though larval collections were made from all aquatic habitat types present at each site identified to contain mosquito larvae. One of our field sites, Belfast Zoo, does not strictly fall under the definition of ‘urban garden’, though trapping here was focused around container-based habitats, thus the site is categorised as ‘urban’ for analyses. Sampling was designed to divide effort equally geographically across the six counties of Northern Ireland, such that one of each habitat type was sampled in at least three counties (Table S1). Sites included nature reserves, privately-owned farmland and private residential gardens.

**Figure 1:**
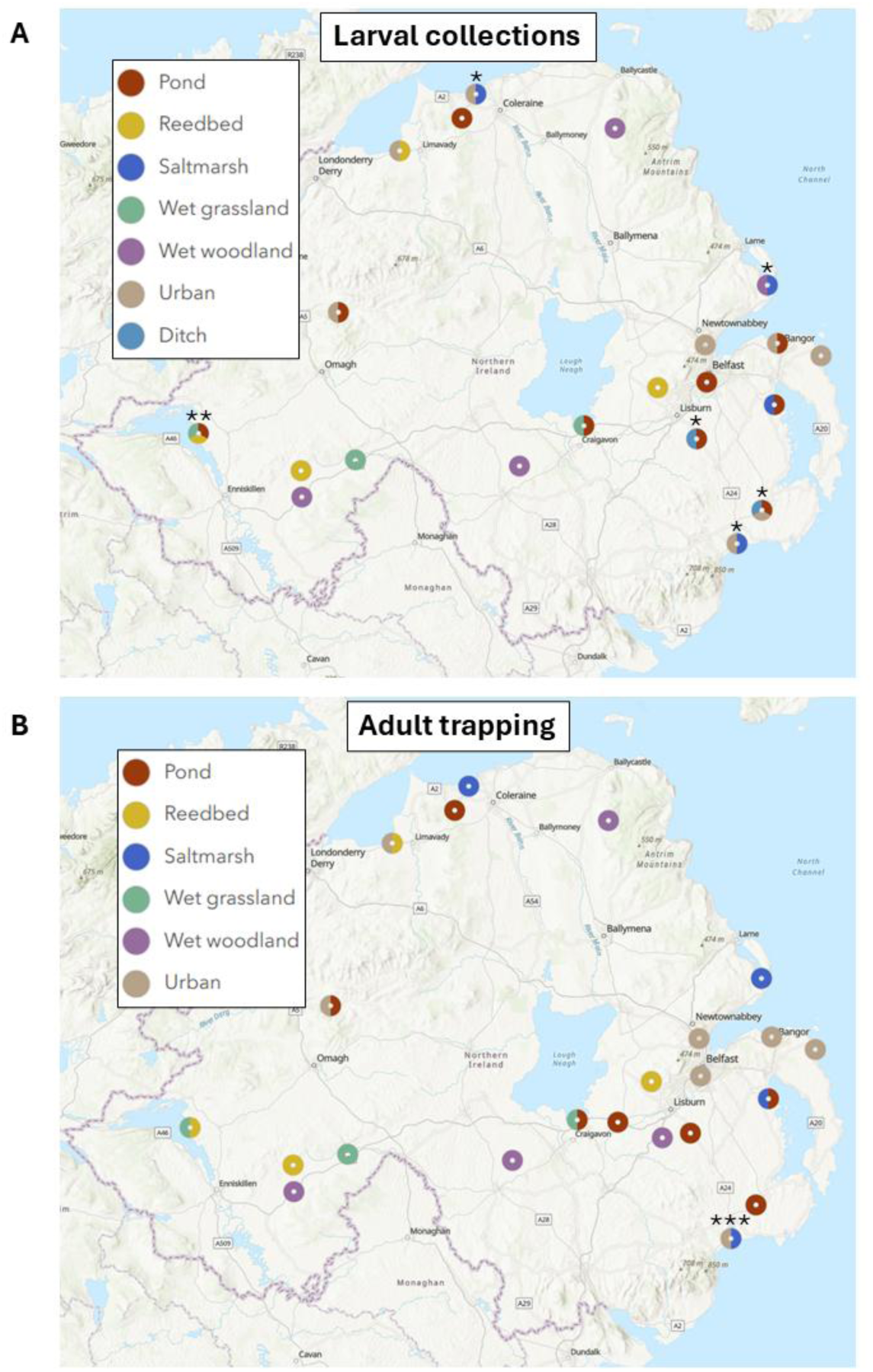
Distribution of sampling sites, coloured by target wetland and organised by survey type (A: larval collections; and B: adult trapping). Sampling sites within 5 kilometres of each other are aggregated for visualisation. Number of colours per circle indicates number of trapping sites, except for circles marked with an asterisk (*) where multiple larval collections were made at a single site, the circle marked with two asterisks (**) where larval collections were made at two sites, one of which contained multiple wetlands (Castle Archdale Country Park; pond and wet grassland) and the circle marked with three asterisks (***) where adult trapping targeted two habitat types in a single site.

Additional ‘one-off’ snapshot surveys were undertaken in July and August 2025 at twelve sites, to opportunistically broaden the coverage of mosquito species collected. Sites were selected based on their close proximity to existing sites and presence of at least one target habitat type. Furthermore, enhanced surveillance (*i.e*., deployment of one additional adult trap at a site, placed at least 500 metres away from the existing trap) was undertaken in July and August 2025 to collect additional mosquitoes at sites noted to contain higher adult abundances (Table S1).

### Larval Sampling

Potential aquatic larval habitats were identified at each site as areas of standing water (*e.g.,* ponds, puddles, ditches, water-filled artificial containers). Three dips were taken at different areas around the periphery of each aquatic habitat using a 300 mL dipper. Dip contents were checked visually for presence of mosquito larvae. Any present mosquito larvae were collected with a pipette and transferred to a falcon tube to be transported for later counting and identification. Collections were made monthly at the same time of trap deployment.

Water pH and temperature of each habitat were recorded using a HI-98128 Water Resistant pH Tester (Hanna Instruments) following the manufacturer’s instructions. Water depth around the periphery of the habitat, approximate habitat size (small, <1 m^2^ in area; medium, 1-3 m^2^ in area; and large, >3 m^2^ in area), vegetation type (none, emergent, floating and submerged) and vegetation coverage (none; low, 1-33% observed coverage; medium, 34-66% observed coverage; and high, 67-100% observed coverage) were recorded. The presence of known mosquito larval predators (*e.g.*, diving beetles or boatmen (Medlock & Snow, 2008)) in the dip was noted.

### Adult Trapping

Adult mosquitoes were sampled using a BG-Pro trap (Biogents), fitted with a BG-Lure attractant and dry ice to generate CO_2_. Traps were deployed within 10 metres of target aquatic habitats on a monthly basis for two consecutive nights at each site, with the exception of Ballycarry in June 2025, where two traps were deployed for just one night (treated as a single collection for statistical analyses and in reporting summaries, to standardise with the 48-hour sampling effort of the other collections). Traps were hung EVS-style on a tree or fence. After two nights, the trap was collected and mosquitoes immediately transferred onto dry ice and stored at -80°C. TGP-4017 Tinytag data loggers (Gemini Data Loggers) were concurrently deployed to record temperature every 30 minutes over the full 48h period. Tags were placed close to the trap, but sufficiently far (at least 50cm away) to prevent dry ice from confounding temperature readings. Data on average daily temperature and total daily rainfall were obtained from the Met Office for the nearest weather stations from each site (Table S1).

### Morphological Identification

Larvae and adults were identified under a microscope using identification keys (Becker et al., 2020) to species/complex level. Early instar larvae were reared in the laboratory until 3^rd^ or 4^th^ instar to facilitate identification. Pupae sampled during larval surveys were reared in the laboratory until adult emergence. When a specimen was morphologically identified as a cryptic taxon (*Culex pipiens*/*torrentium* or *Anopheles maculipennis* s.l. (Falleroni)), legs or other suitable tissue were removed and stored in 100% ethanol for subsequent molecular identification. All specimens of *Anopheles maculipennis* s.l., and all adult specimens of *Culex pipiens/torrentium* were identified to species-level. For larval samples of *Culex pipiens/torrentium* where the total catch exceeded 30, a subset of 30 was randomly selected to be identified to species-level.

### DNA Extraction & Molecular Identification

DNA was extracted from either whole-body (in the case of larvae), legs or wings (in the case of adults) from specimens selected for molecular identification. DNA was extracted through grinding the tissue with a pipette tip containing 20/100µL (legs or wings/whole-body) of buffer consisting of a mix of 10 mM Tris HCl, 1 mM EDTA, 25 mM NaCl, and 10 µL proteinase K (added fresh). This solution was incubated at 37°C for 20 minutes, before proteinase K was inactivated by heating to 95°C for 2 minutes.

Extracted DNA belonging to *Culex pipiens/torrentium* specimens was used to identify species/biotype by PCR using the primers and following the protocols of (Bahnck & Fonseca, (2006) and (Smith & Fonseca, (2004), while specimens of *Anopheles maculipennis* s.l. were tested using the primers of (Proft et al., (1999) and (Kronefeld et al., (2014). Primer sequences are shown in Table S2. PCR products were analysed using gel electrophoresis on a 2.5% agarose gel, and results visualised using the Biorad GelDoc Go Imaging System.

### Data Analysis

Statistical analyses were carried out in R v4.4.2 (R Core Team, 2024) using R Studio (Posit team, 2026). Analyses were focussed on two outcome variables of interest: mosquito presence (yes/no; larvae only), and mosquito abundance (larvae and adults separately). Larval mosquito abundance was the number of individuals collected across the three dips, while adult mosquito abundance was the number of mosquitoes collected in a single trap (‘snapshot’ and ‘enhanced’ collections were treated as independent datapoints). The relationship between mosquito response variables and environmental data was analysed using generalised linear mixed effects models with the R package glmmTMB (Brooks et al., 2017; McGillycuddy et al., 2025). Larval and adult stage abundance data were analysed separately. We first fit a model with all species pooled per life stage, with separate analyses secondly performed for potential vector or nuisance species, *Culex pipiens* s.l. and *Culiseta annulata* (adult stage only). The presence of *Culex* spp. larvae was also analysed in a separate model. A total of six models were fit in the present study, and initial models are summarised in Table 1.

**Table 1:**
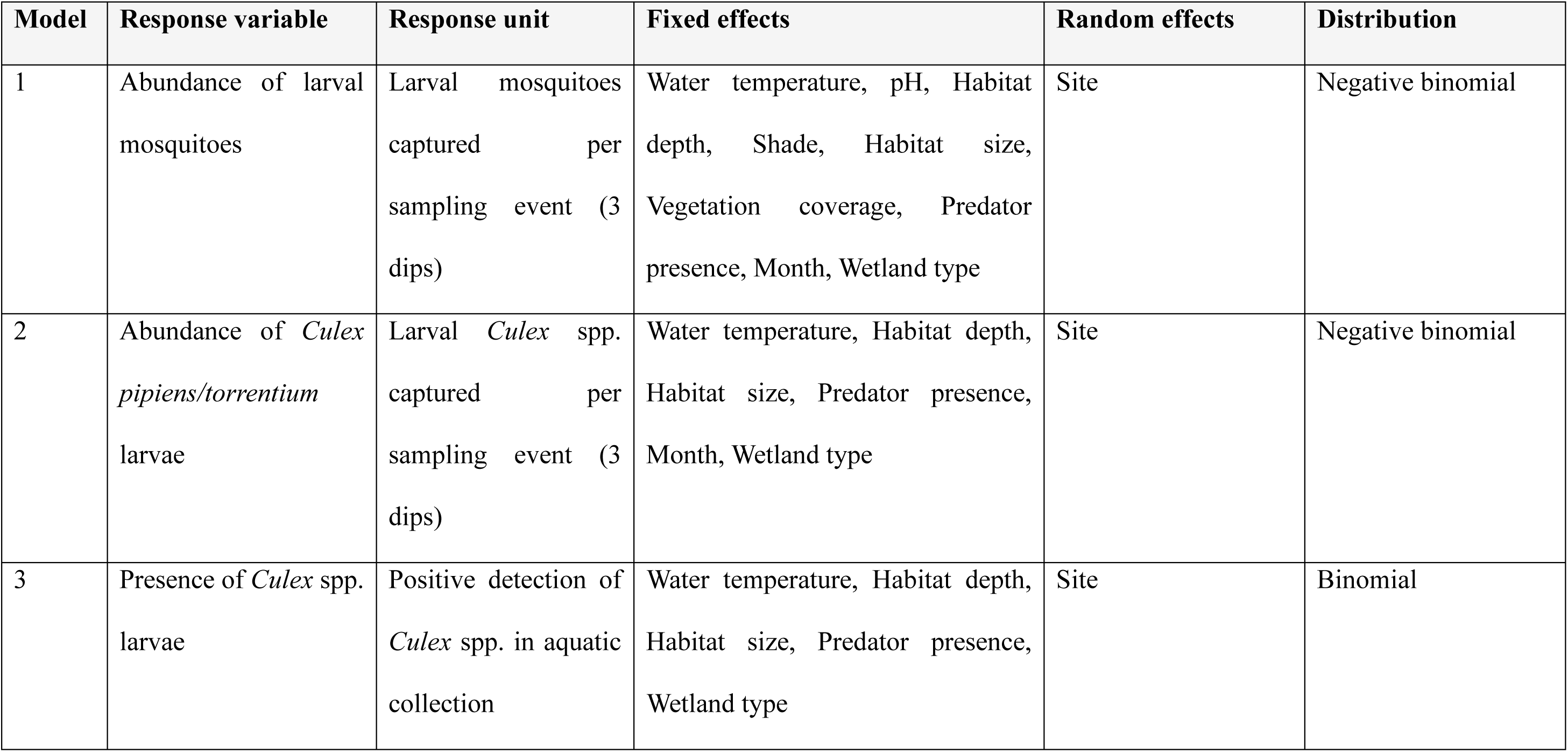
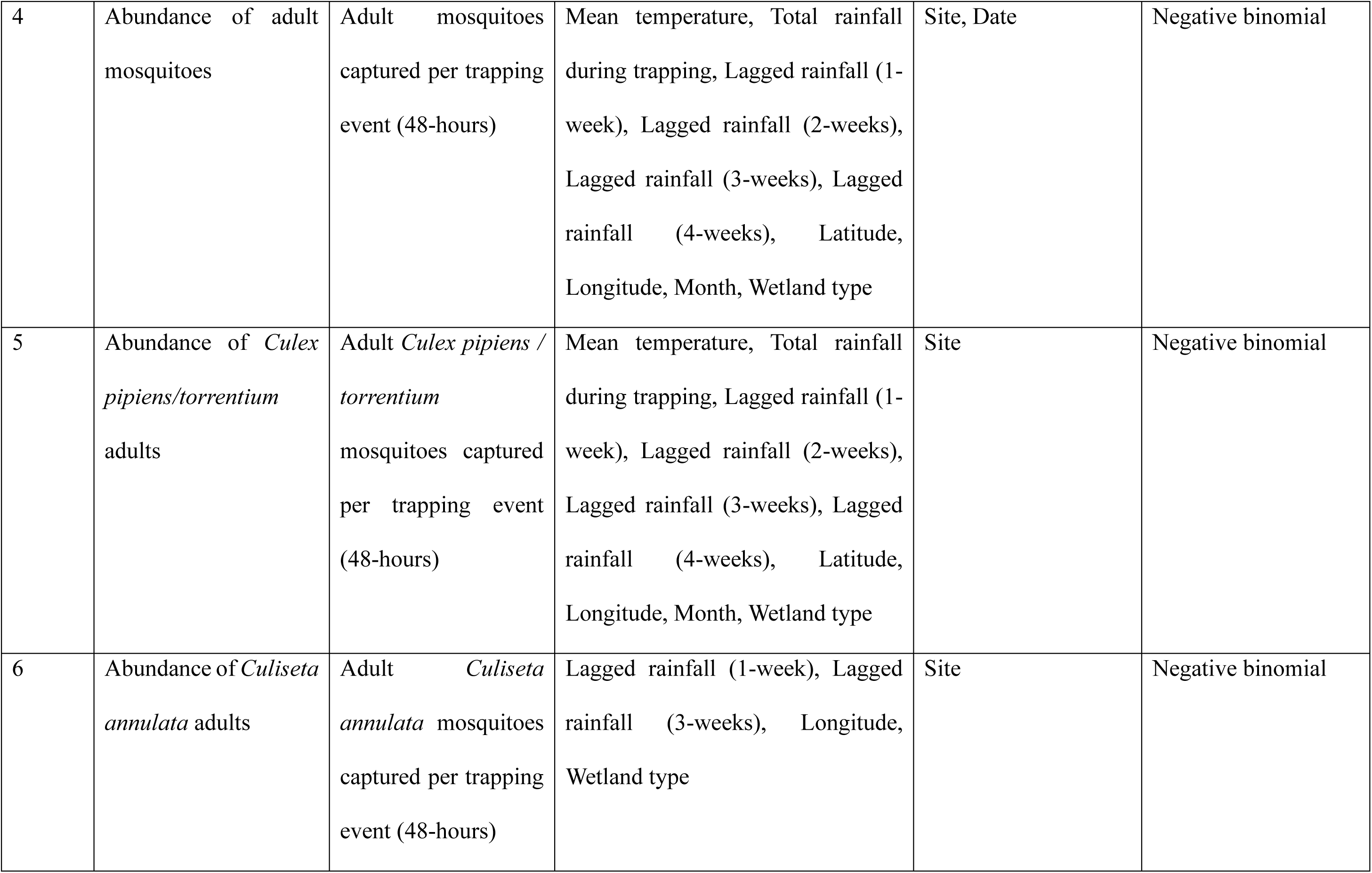
Summary of GLMM statistical models tested in this study. All mosquito abundance models were fit with a negative binomial distribution due to the high number of zeroes in the dataset, whilst the *Culex* larval stage presence model was fit with a binomial distribution.

Explanatory variables (Table 1) were checked for collinearity by computing the variance inflation factor using the R package car (Fox & Weisberg, 2019), considering a threshold value of 5. Explanatory variables tested were water temperature (snapshot), pH, habitat depth, shade, habitat size, vegetation coverage, predator presence, month and wetland type (larval analyses); mean temperature, rainfall during trapping (mm), lagged rainfall (mm; 1, 2, 3 and 4-weeks), latitude, longitude, month and wetland type (adult analyses). Site was treated as a random effect and sampling date was included as a random effect where model convergence was possible (model 4 only). Temperature values used in statistical models of adults were mean values (over the 48-hour sampling period) as obtained from the Tinytag data logger, and where these data were unavailable, the mean temperature during the trapping period was obtained from the nearest weather station (17/98 trapping events). Initial models were checked for overdispersion, zero inflation and goodness of fit using residual simulations through the R package DHARMa (Hartig, 2024). Poisson distributions were initially used for abundance data, though residual diagnostics found this to be unsuitable due to zero-inflation, and so the negative binomial distribution was implemented. All continuous explanatory data were centred and scaled to allow for model convergence. Nested models were compared using likelihood ratio tests, with stepwise backward elimination used to determine the simplest statistically significant model. Due to the high number of zeros in our dataset, formulated models could not converge with all explanatory variables when moving from all-species models to species-specific models. Thus, a reduced number of explanatory variables was incorporated into species-specific models to achieve model fit (Table 1). Post-hoc Tukey tests were conducted on significant categorical variables using the R package emmeans (Lenth & Piaskowski, 2025). Predicted relationships were extracted from final models using the R package ggeffects (Lüdecke, 2018) and visualised using ggplot2 (Wickham, 2016).

## Results

### Wetland microclimates

The mean recorded temperature during the study period was 13.99°C, and mean temperatures and total rainfall are summarised for each weather station in Table S3 and visualised in Figure S1. Mean temperatures during the study period were generally higher than the 1991-2020 Met Office average (Table S4).

### Ecology of larval stage mosquitoes

Mosquito larvae were detected in 41 of 76 sampling events; corresponding to a site positivity of 68% (19/28). Sampling effort (*i.e.*, number of sampling events) was spread across May (n = 12), June (n = 16), July (n = 23) and August (n = 25). Sampling effort was greater in July and August due to snapshot surveys and identification of additional aquatic habitats at sites. A total of 1479 larvae were collected in the present study (Table 2, Table S5) representing seven species across four genera: *Culex pipiens*, *Culex torrentium*, *Aedes detritus*, *Culiseta annulata*, *Culiseta morsitans*, *Anopheles claviger* and *Anopheles maculipennis* s.l. Larval specimens from *Anopheles maculipennis* s.l. could not be identified to species-level. Potential mosquito predators were detected in 29 of 76 collections. No attempt was made to identify these to species level, but predator taxa detected included Anisoptera (n = 1), Anura (n = 2), Corixidae (n = 5), Dytiscidae (n = 13), Gammaridae (n = 9) and Zygoptera (n = 1). Typically, only one predatory taxon was observed per sampling event, except for one occasion where Anura and Dytiscidae were detected together in the same habitat.

**Table 2:** Sampling and habitat association data by mosquito species caught as larvae. Numbers in parentheses indicates the total number of collections where a given species was detected out of the total number of collections made per habitat. Here, *Culex pipiens* s.l. includes *Culex pipiens pipiens*, *Culex pipiens/molestus* hybrids and *Culex torrentium*, as not every larval specimen of *Culex* was tested by PCR to species/biotype level.

| Species | Total abundance (number of collections with species detected/total number of collections per habitat) |  |  |  |  |  |  |  |
| --- | --- | --- | --- | --- | --- | --- | --- | --- |
|  | Artificial<br>container<br>(14) | Ditch (4) | Pond (17) | Reedbed<br>(14) | Saltmarsh<br>(12) | Wet<br>grassland<br>(7) | Wet<br>woodland<br>(8) | Total |
| *Culex pipiens* s.l. | 233<br>(10/14) | 271 (3/4) | 370 (3/17) | 219 (5/14) | 4 (1/12) | 15 (2/7) | 27 (1/8) | 1139 (25/76) |
| *Aedes detritus*<br>(Haliday) | 0 (0/14) | 0 (0/4) | 0 (0/17) | 0 (0/14) | 279 (9/12) | 0 (0/7) | 0 (0/8) | 279 (9/76) |
| *Culiseta annulata*<br>(Schrank) | 0 (0/14) | 0 (0/4) | 11 (2/17) | 31 (3/14) | 0 (0/12) | 0 (0/7) | 5 (3/8) | 47 (8/76) |
| *Culiseta morsitans*<br>(Theobald) | 0 (0/14) | 9 (1/4) | 0 (0/17) | 0 (0/14) | 0 (0/12) | 0 (0/7) | 0 (0/8) | 9 (1/76) |
| *Anopheles claviger*<br>(Meigen) | 0 (0/14) | 0 (0/4) | 3 (2/17) | 0 (0/14) | 0 (0/12) | 0 (0/7) | 0 (0/8) | 3 (2/76) |
| *Anopheles maculipennis* s.l. | 0 (0/14) | 0 (0/4) | 2 (2/17) | 0 (0/14) | 0 (0/12) | 0 (0/7) | 0 (0/8) | 2 (2/76) |
| **Total mosquitoes** | **233** | **280** | **386** | **250** | **283** | **15** | **32** | **1479** |

Only significant associations are reported here, though all statistical results to identify ecological drivers of mosquito presence and abundance are summarised in Table S6. Larval mosquito abundance was positively associated with low vegetation coverage (χ^2^ = 21.046, df = 3, p < 0.001; Figure 2A) and predator absence (χ^2^ = 8.754, df = 1, p < 0.01; Figure 2B). Post-hoc analyses confirmed that larval abundance was greater in the absence of vegetation relative to all other vegetation coverage types (p < 0.05). Larval mosquito abundance differed significantly among wetland types (χ^2^ = 19.140, df = 6, p < 0.01; Figure 2C). Saltmarsh pools had a greater abundance of mosquitoes than all other habitat types (p < 0.05) except for reedbeds and temporary ditches. Artificial containers had a significantly lower abundance of larvae relative to both reedbeds (p < 0.05) and ditches (p < 0.01).

**Figure 2:**
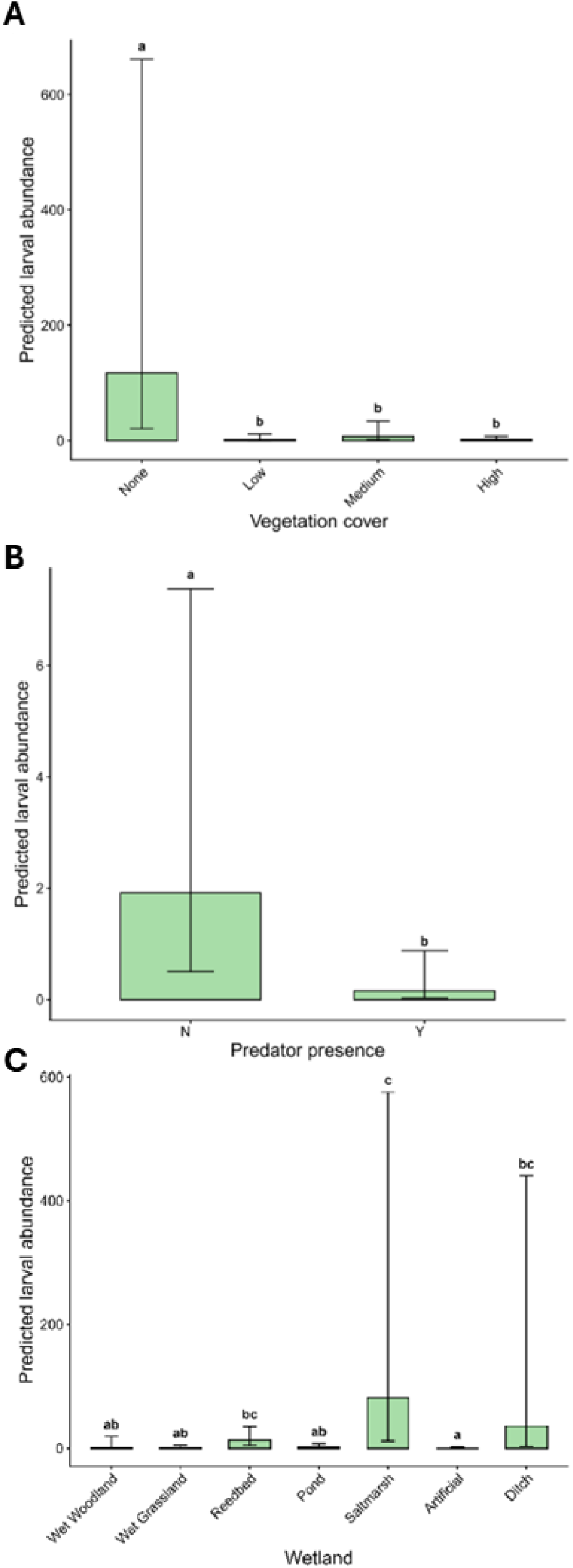
Predicted relationships between abundance of larval mosquitoes per collection and environmental conditions found to be statistically significant. Error bars indicate 95% confidence intervals. (A) Relationship between predicted larval abundance and vegetation cover; (B) Relationship between predicted larval abundance and predator presence; and (C) Relationship between predicted larval abundance and wetland type.

The presence and abundance of vector species *Culex pipiens/torrentium* larvae was next analysed with respect to environmental and ecological drivers. The probability of detecting *Culex pipiens/torrentium* larvae increased with water temperature (χ^2^ = 17.625, df = 1, p < 0.01; Figure 3A) and in the absence of predators (χ^2^ = 11.345, df = 1, p < 0.001; Figure 3B), while differing by wetland type (χ^2^ = 5.841, df = 6, p = 0.02; Figure 3C). Relative to artificial containers, saltmarsh pools (p < 0.05) and ponds (p < 0.05) were significantly less likely to have *Culex* larvae present, though post-hoc analysis did not reveal significant pairwise differences.

**Figure 3:**
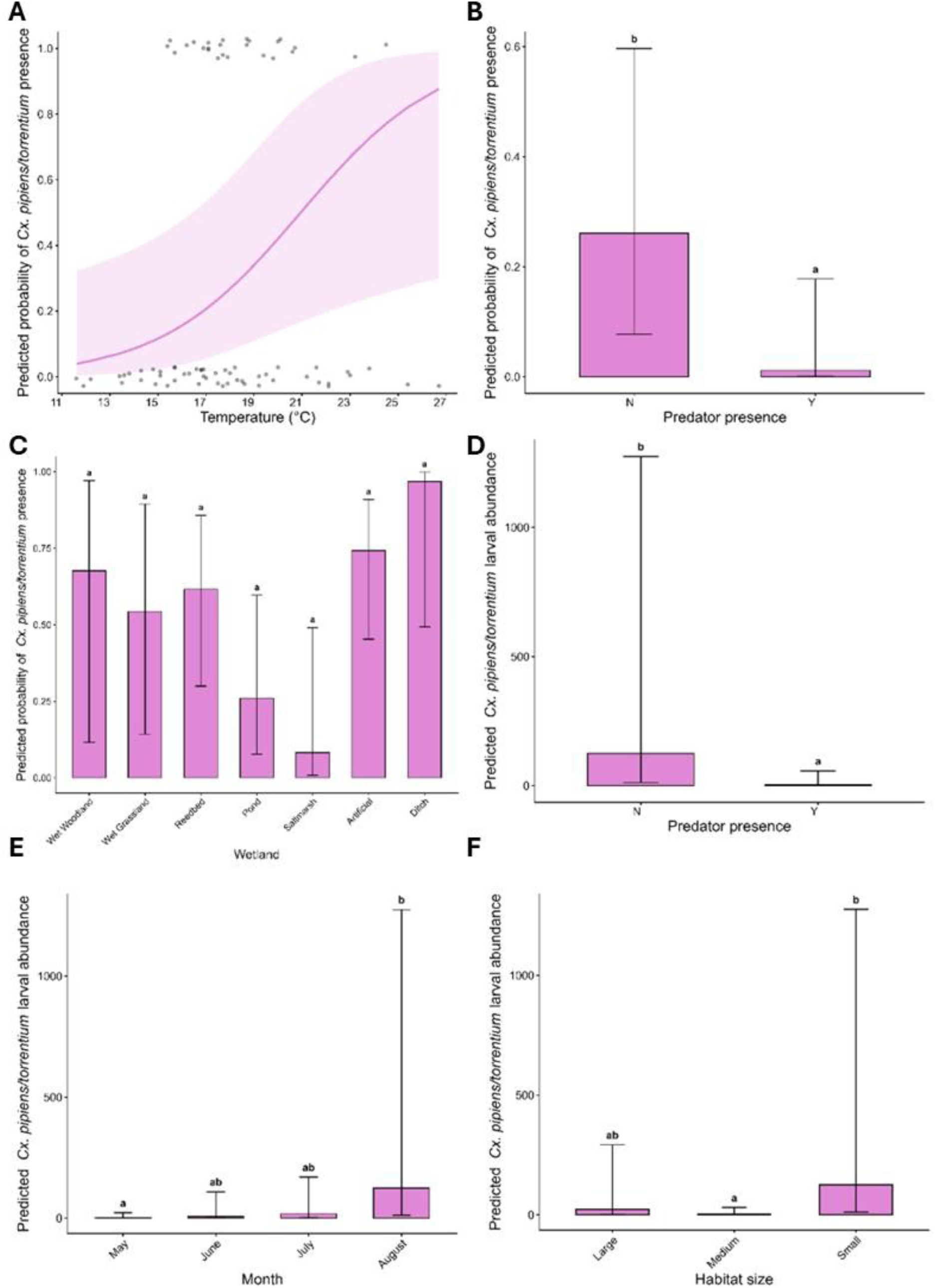
Predicted relationships between larval stage *Culex pipiens/torrentium* presence/abundance and environmental conditions found to be statistically significant. Shaded areas and error bars indicate 95% confidence intervals. (A) Relationship between presence of larval *Culex pipiens/torrentium* and water temperature (°C); (B) Relationship between presence of larval *Culex pipiens/torrentium* and detectable predators present; (C) Relationship between presence of larval *Culex pipiens/torrentium* and wetland type; (D) Relationship between abundance of larval *Culex pipiens/torrentium* and detectable predators present; (E) Relationship between abundance of larval *Culex pipiens/torrentium* and month; and (F) Relationship between abundance of larval *Culex pipiens/torrentium* and habitat size.

*Culex pipiens/torrentium* larval abundance was significantly lower in the presence of predators (χ^2^ = 7.727, df = 1, p < 0.01; Figure 3D), differed among months (χ^2^ = 9.837, df = 3, p = 0.02; Figure 3E) and by habitat size (χ^2^ = 7.292, df = 2, p = 0.03; Figure 3F). *Culex* larvae occurred at a greater abundance in August relative to May (p < 0.05), though no significant differences were detected among other months. Small habitats had significantly more *Culex* larvae than medium-sized habitats (p < 0.05).

### Ecology of adult stage mosquitoes

Adult mosquitoes were detected in 70 of 98 trap collections, representing 23 of the 28 sites. This yielded a total of 951 adult mosquitoes, representing fifteen species across five genera (Figure 4, Table 3) and representing all eight functional groups (Figure S2). *Culex pipiens pipiens* was the most abundant species detected (n = 444) and was frequently detected and in various sites, appearing in 51 out of 98 sampling events. *Culiseta annulata* (n = 172) and *Aedes detritus* (n = 85) were also abundant and frequently detected. Notably, *Aedes detritus*, the saltmarsh mosquito, was detected on several occasions in Ballykelly ICW (reedbed) and Ballykelly Village (urban garden), both sites of which were located approximately 3 kilometres from the nearest saltmarsh site. All adult specimens from *Anopheles maculipennis* s.l. (n = 10) were identified to species level by PCR and identified as *Anopheles messeae*, though no result could be obtained on larval specimens.

**Figure 4:**
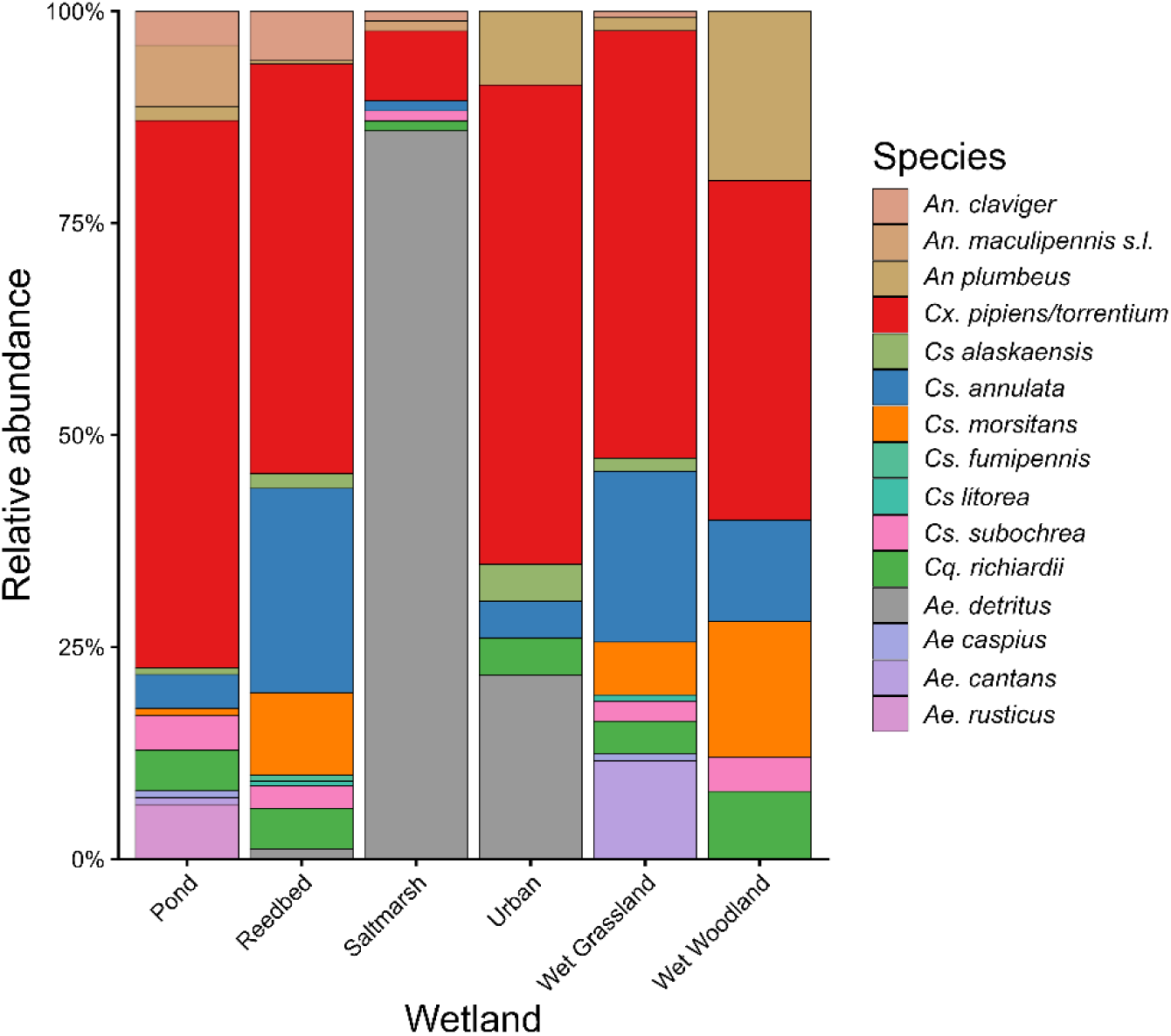
Relative abundance of adult mosquito species collected in Northern Ireland by wetland type sampled.

**Table 3:** Sampling and habitat association data by mosquito species caught as adults. Numbers in parentheses indicates the total number of collections where a given species was detected out of the total number of collections made per habitat. Here, *Culex pipiens* (Linnaeus) includes all specimens identified by morphology as *Culex pipiens* s.l. and identified by PCR as either *Culex pipiens pipiens*, *Culex pipiens/molestus* hybrids or not determined (*i.e.* negative PCR result). *Denotes the first detection of this species on the island of Ireland.

| Species | Total abundance (number of collections with species detected/total number of collections per habitat) |  |  |  |  |  |  |
| --- | --- | --- | --- | --- | --- | --- | --- |
|  | Pond | Reedbed | Saltmarsh | Urban | Wet<br>grassland | Wet<br>woodland | Total |
| *Culex pipiens* (Linnaeus) | 80 (14/20) | 273 (14/17) | 7 (4/15) | 11 (6/16) | 64 (11/15) | 9 (2/15) | 444 (51/98) |
| *Culiseta annulata*<br>(Schrank) | 5 (4/20) | 136 (11/17) | 1 (1/15) | 1 (1/16) | 26 (9/15) | 3 (3/15) | 172 (29/98) |
| *Aedes detritus* (Haliday) | 0 (0/20) | 7 (3/17) | 73 (9/15) | 5 (3/16) | 0 (0/15) | 0 (0/15) | 85 (15/98) |
| *Culiseta morsitans*<br>(Theobald) | 1 (1/20) | 55 (8/17) | 0 (0/15) | 0 (0/16) | 8 (4/15) | 4 (1/15) | 68 (14/98) |
| *Coquellittidia richiardii*<br>(Ficalbi) | 6 (3/20) | 27 (5/17) | 1 (1/15) | 1 (1/16) | 5 (3/15) | 2 (2/15) | 42 (15/98) |
| *Anopheles claviger*<br>(Meigen) | 5 (3/20) | 33 (8/17) | 1 (1/15) | 0 (0/16) | 1 (1/15) | 0 (0/15) | 40 (13/98) |
| *Culiseta subochrea*<br>(Edwards) | 5 (2/20) | 15 (5/17) | 1 (1/15) | 0 (0/16) | 3 (1/15) | 1 (1/15) | 25 (10/98) |
| *Aedes cantans* (Meigen) | 1 (1/20) | 0 (0/17) | 0 (0/15) | 0 (0/16) | 15 (2/15) | 0 (0/15) | 16 (3/98) |
| *Culiseta alaskaensis*<br>(Felt) | 1 (1/20) | 10 (3/17) | 0 (0/15) | 1 (1/16) | 2 (1/15) | 0 (0/15) | 14 (6/98) |
| *Anopheles plumbeus*<br>(Stephens) | 2 (1/20) | 2 (1/17) | 0 (0/15) | 2 (1/16) | 2 (1/15) | 5 (2/15) | 13 (6/98) |
| *Anopheles messeae* *<br>(Falleroni) | 9 (3/20) | 0 (0/17) | 1 (1/15) | 0 (0/16) | 0 (0/15) | 0 (0/15) | 10 (4/98) |
| *Aedes rusticus* (Rossi) | 8 (3/20) | 0 (0/17) | 0 (0/15) | 0 (0/16) | 0 (0/15) | 0 (0/15) | 8 (3/98) |
| *Culiseta litorea* (Shute) | 0 (0/20) | 3 (3/17) | 0 (0/15) | 0 (0/16) | 1 (1/15) | 0 (0/15) | 4 (4/98) |
| *Culiseta fumipennis* *<br>(Stephens) | 0 (0/20) | 4 (2/17) | 0 (0/15) | 0 (0/16) | 0 (0/15) | 0 (0/15) | 4 (2/98) |
| *Culex torrentium*<br>(Martini) | 0 (0/20) | 0 (0/17) | 0 (0/15) | 2 (2/16) | 1 (1/15) | 1 (1/15) | 4 (4/98) |
| *Aedes caspius* (Pallas) | 1 (1/20) | 0 (0/17) | 0 (0/15) | 0 (0/16) | 1 (1/15) | 0 (0/15) | 2 (2/98) |
| **Total mosquitoes** | **124** | **565** | **85** | **23** | **129** | **25** | **951** |

Adult mosquito abundance was negatively associated with rainfall 1 week prior (χ^2^ = 3.976, df = 1, p < 0.05; Figure 5A), and differed among months (χ^2^ = 19.847, df = 3, p < 0.001; Figure 5B) and wetland types (χ^2^ = 26.117, df = 5, p < 0.0001; Figure 5C). May had a significantly lower abundance of adult mosquitoes relative to all months (p < 0.05), with no significant variation between other months. Both reedbeds and wet grasslands had significantly higher adult abundance than urban habitats (both: p < 0.05). Wet woodlands had a significantly lower abundance of adult mosquitoes relative to ponds (p < 0.05), reedbeds (p < 0.0001) and wet grasslands (p < 0.05).

**Figure 5:**
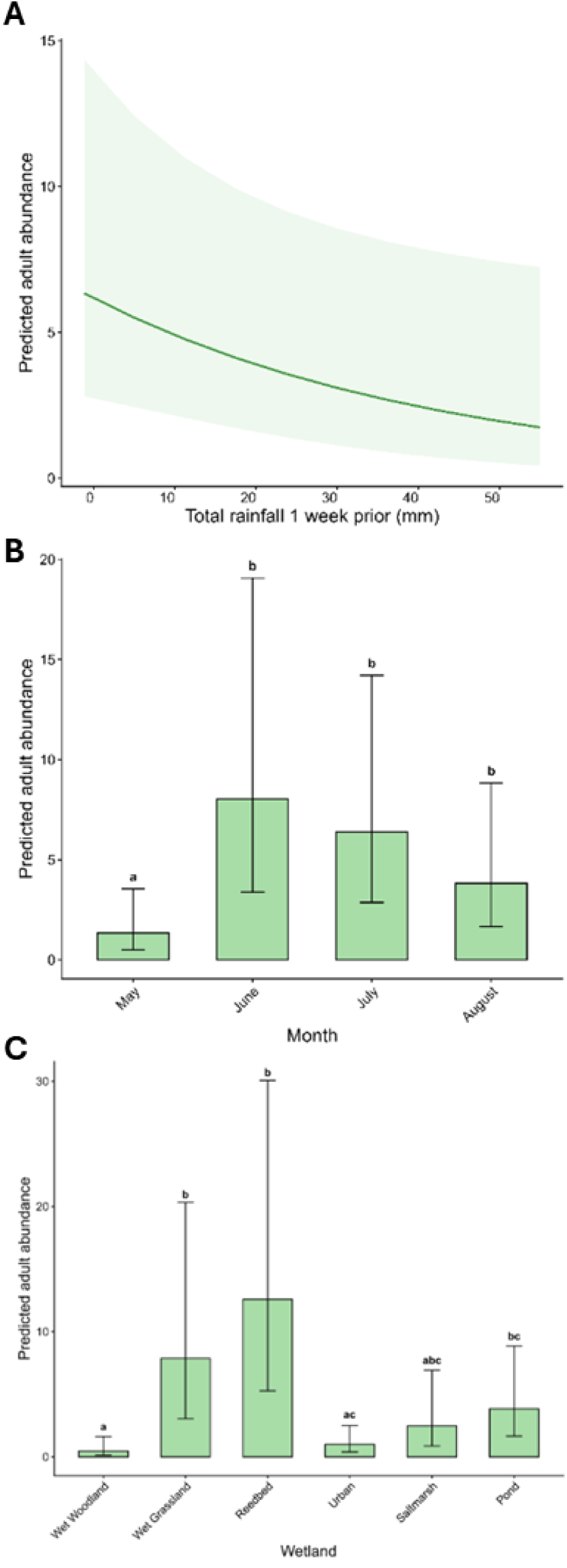
Predicted relationships between total number of adult mosquitoes and environmental conditions found to be statistically significant. Shaded areas and error bars indicate 95% confidence intervals. (A) Relationship between predicted adult abundance and lagged total rainfall 1 week prior (mm); (B) Relationship between predicted adult abundance and month; and (C) Relationship between predicted adult abundance and wetland type.

Abundances of both potential vector species *Culex pipiens/torrentium* and *Culiseta annulata* were next analysed with respect to environmental and ecological drivers. Adult *Culex pipiens/torrentium* mosquitoes were negatively correlated with rainfall 1-week prior (χ^2^ = 4.221, df = 1, p < 0.05; Figure 6A), but positively correlated with rainfall 4-weeks prior (χ^2^ = 3.843, df = 1, p < 0.05; Figure 6B). Adult *Culex pipiens/torrentium* were also significantly more abundant in southerly latitudes (χ^2^ = 5.867, df = 1, p < 0.05; Figure 6C), and differed among months (χ^2^ = 19.847, df = 3, p < 0.01; Figure 6D) and wetland types (χ^2^ = 25.748, df = 3, p < 0.0001; Figure 6E). *Culex* adult abundance was lower in May than other sampling months (p < 0.05), and higher in reedbeds relative to all other wetland types (p < 0.05), except for ponds and wet grasslands.

**Figure 6:**
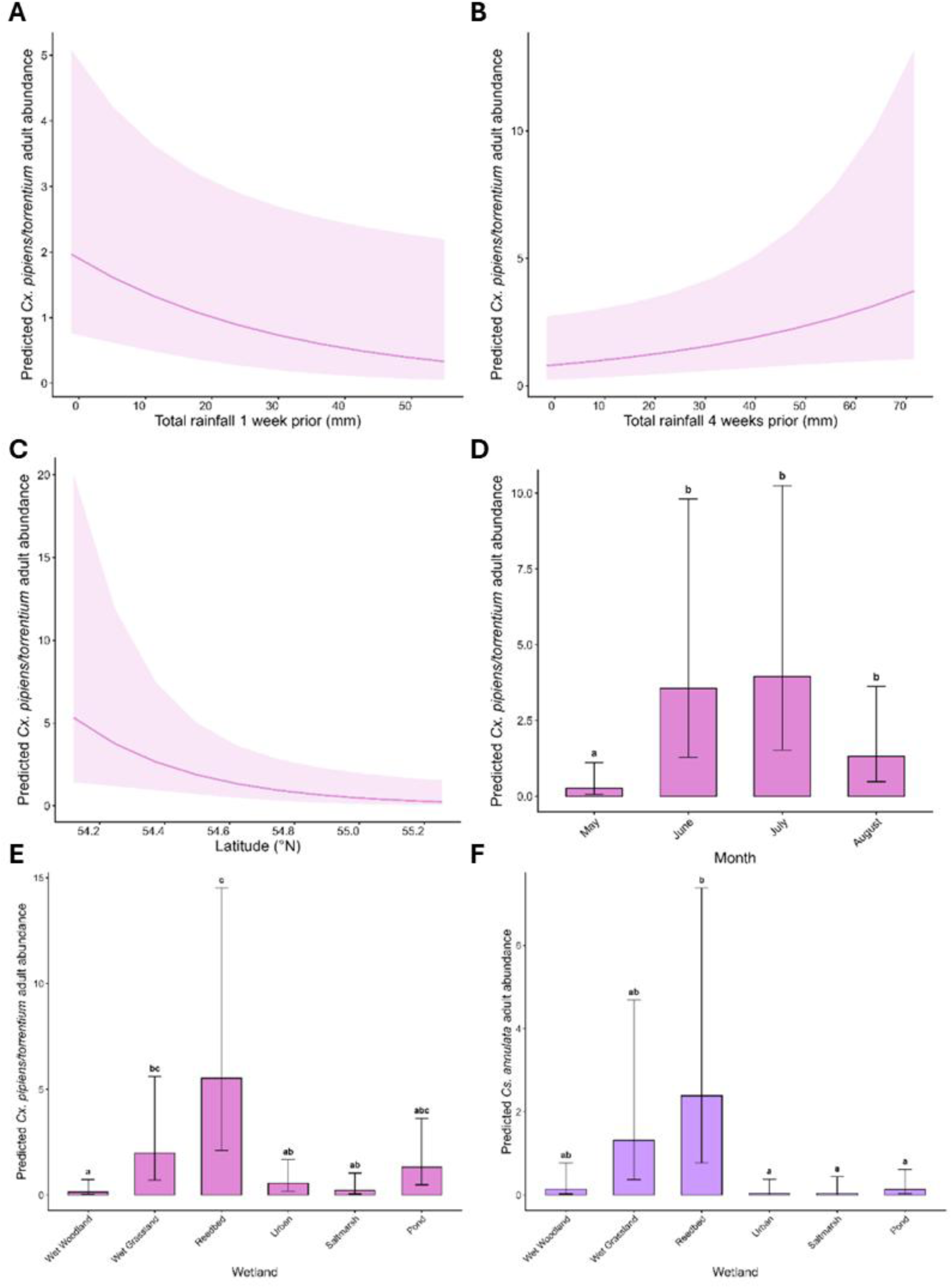
Predicted relationships between individual mosquito species (adult stage) and environmental conditions found to be statistically significant. Shaded areas and error bars indicate 95% confidence intervals. (A) Relationship between adult *Culex pipiens/torrentium* and lagged rainfall 1 week prior (mm); (B) Relationship between adult *Culex pipiens/torrentium* and lagged rainfall 4 weeks prior (mm); (C) Relationship between adult *Culex pipiens/torrentium* and latitude; (D) Relationship between adult *Culex pipiens/torrentium* and month; (E) Relationship between adult *Culex pipiens/torrentium* and wetland type; and (F) Relationship between adult *Culiseta annulata* and wetland type.

Adult *Culiseta annulata* mosquito abundance also differed among wetland types (χ^2^ = 21.219, df = 1, p < 0.001; Figure 6F), being higher in reedbeds than ponds, saltmarsh and urban habitats (p < 0.05). No other significant environmental predictors of *Cs. annulata* were identified.

### Distribution of *Culex* mosquito species/biotypes

A subset of 550 field-collected *Culex* mosquito larvae was selected for molecular analysis to identify species/biotype, from which 294 (53%) could be identified. All 444 field-collected *Culex* mosquito adults were tested by PCR to identify species/biotype, with positive identification obtained for 426 specimens (96%). In combination, this gave a total sample size of 720 *Culex* samples identified to species/biotype level. Differences in extracted tissue between adult and larval mosquitoes may explain the variation in identification success rate (Text S1).

Most *Culex* specimens were identified as *Cx. p. pipiens* (n = 643; 89.3%), followed by *Cx. torrentium* (n = 67; 9.3%). Evidence of potential hybridisation between *Cx. p. pipiens* and *Cx. p. molestus* was uncovered (n = 13; 1.8%) though no specimens were identified as *Cx. p. molestus*. *Culex. p. pipiens* had a widespread distribution across Northern Ireland (detected in twenty-three sites), with adults detected in all habitat types though less frequently in wet woodlands (Figure 7A). Hybrids were collected at five sites in total, one reedbed where adults were collected in June and August, and four others in the southern half of the country in different habitats (urban garden, saltmarsh pools and ponds). Most detections of hybrids (n = 7/13; 53.85%) occurred in agricultural settings. *Culex torrentium* was detected at nine sites, and was more frequently detected in larval collections than in adult traps (Figure 7B). In artificial containers, *Cx. torrentium* occurred at a greater density relative to other larval habitats (Figure 7C).

**Figure 7:**
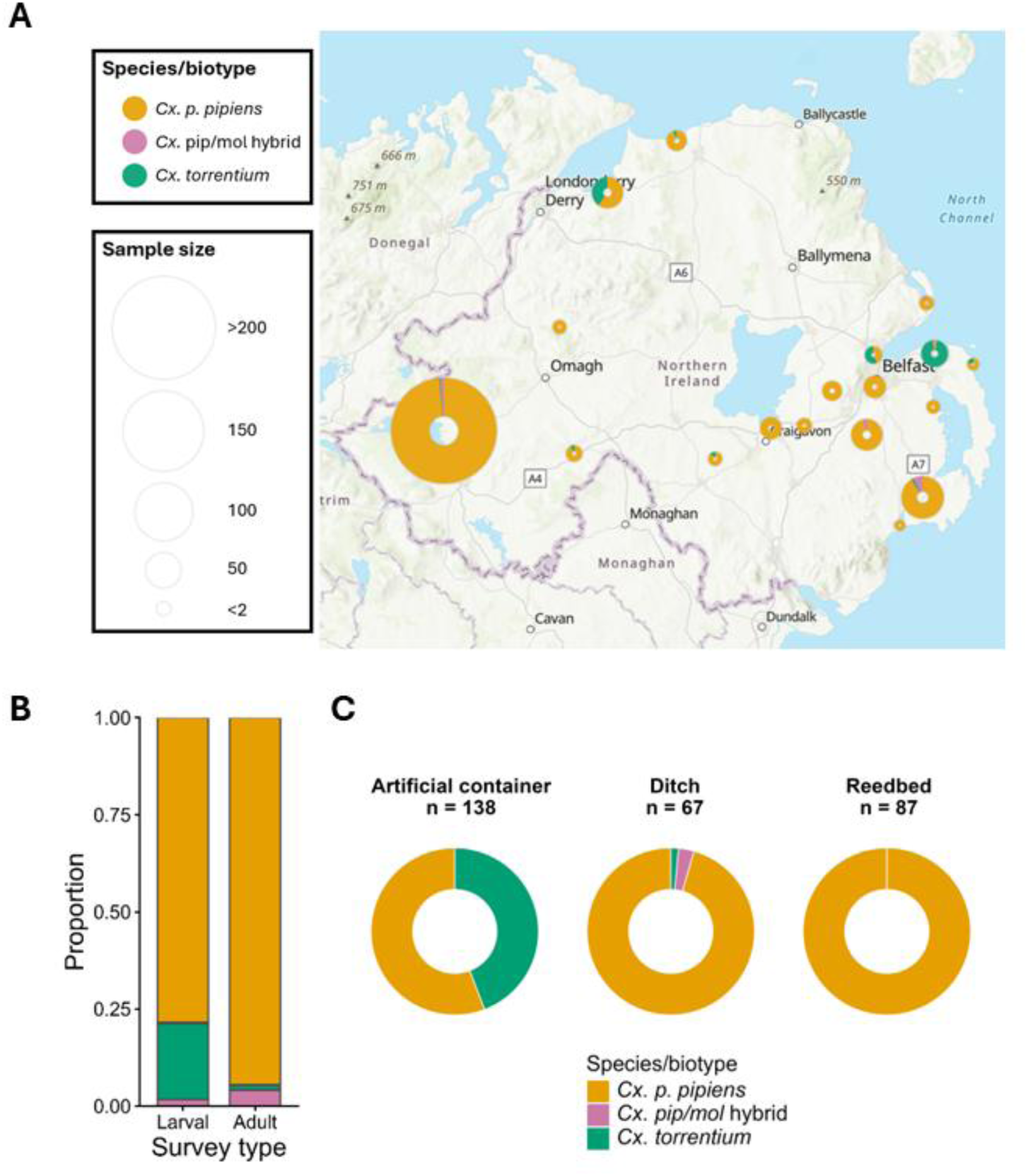
Relative abundance and distribution of *Culex* mosquito species/biotypes across Northern Ireland. (A) Map visualising geographic distribution of *Culex* species/biotypes identified by both adult and larval surveys. Sites within 5 kilometres are grouped together for visualisation. (B) Relative proportion of *Culex* species/biotypes according to larval/adult survey type. (C) Relative proportion of *Culex* species/biotypes in the top 3 most frequently occupied larval habitats.

## Discussion

Here, we report results from the first nationwide survey of mosquitoes in Northern Ireland, investigating six key habitat types selected to provide a comprehensive overview of resident mosquito species. Mosquitoes had a widespread distribution across Northern Ireland, being detected at almost every site surveyed (26/30). Potential vector species, *e.g.*, *Culex pipiens pipiens* and *Culex torrentium*, were widely distributed, and the potential human-avian reservoir bridge vector *Culex pipiens/molestus* hybrids were confirmed for the first time on the island of Ireland. Additionally, *Culiseta annulata* and *Aedes detritus*, each of which are understood to be potential vectors of West Nile and/or Usutu viruses, were found in high abundances (Fros et al., 2015; Pilgrim et al., 2024). Thus, we emphasise the need for continued and enhanced surveillance of both mosquitoes and mosquito-borne pathogens in Northern Ireland to better understand the risk to human and wildlife health in this data-poor context.

Fifteen of the twenty-one previously reported mosquito species on the island of Ireland (Schaffner et al., 2024) were found in the present study, in addition to first records of *Culex pipiens/molestus* hybrid biotype, *Culiseta fumipennis* and *Anopheles messeae;* this brings the total number of recorded mosquito fauna in Ireland to twenty-three. Thirty-six mosquito species (and all of the species represented here) are native to Great Britain (*i.e*. England, Scotland and Wales; Harbach et al., 2017), with Irish mosquito fauna representing a subset of these. *Culiseta fumipennis* is a lesser known species, with few details known on its phenology or feeding behaviour, though it has never been documented to bite humans (Becker et al., 2020), and so it likely poses little risk as a zoonotic vector. *An. messeae* belongs to the *Anopheles maculipennis* complex and is particularly closely related to *An. daciae*, which was recently detected in the Republic of Ireland (Schaffner et al., 2024). This species is closely associated with agricultural land uses and has been documented to bite both humans and livestock (Fyodorova et al., 2006), potentially enabling it to act as a localised nuisance biter, though it was not detected at high densities here. Both *Cs. fumipennis* and *An. messeae* have previously been detected in Great Britain, and there may be additional species as yet undiscovered on the island of Ireland. Additional details on the natural history of species not collected here are provided in Text S2. Continued surveillance will be important to uncovering the potential presence of additional mosquito species in Northern Ireland, as well as to monitor for potential incursion of non-native taxa.

The widespread abundance of *Culex* mosquitoes recorded across Northern Ireland here is expected. Both species (*Cx. pipiens* and *Cx. torrentium*) are common and abundant through Europe, and known to colonise both natural and man-made habitats (Krol et al., 2024). Given the primary role of *Cx. pipiens* in transmitting emerging arboviruses throughout Europe (*e.g.*, Jansen et al., 2023; Krambrich et al., 2024; Soto et al., 2023; Vogels et al., 2017), understanding the environmental drivers of its abundance both locally and continentally is important to assess epidemiological risk. The abundance of *Culex* larvae was lower in the presence of known predators, suggesting that natural enemies could play an important role in regulating larval populations. Interestingly, this effect was not detected in another study of *Culex* in England (Golding et al., 2015). The abundance of adult *Cx. pipiens* had a strong negative association with lagged rainfall at 1 week, and a positive association with lagged rainfall at 4 weeks. Adult activity may have been discouraged by recent (preceding 1 week) rainfall, *e.g.,* in the United States, it has been demonstrated that rainfall has a negative association with adult longevity (Jones et al., 2012) and could potentially flush out larvae and pupae (Gardner et al., 2012), though this effect is likely dependent on rainfall intensity. *Culex pipiens* is a multivoltine species in Great Britain (Ewing et al., 2019), with Northern populations estimated to have a median developmental duration of around 3 weeks at 18°C (Parvy et al., 2026), and thus lagged rainfall from 4 weeks could have created suitable habitat (*e.g.,* by accumulation of water in ditches or artificial containers) for a previous generation to oviposit, larvae to hatch, develop and emerge into blood-seeking adults. Overall, the abundance of both larval and adult stages of *Cx. pipiens* is closely associated with environmental and ecological drivers, which will be key to assessing future risk of *Culex*-borne arboviruses.

Of immediate relevance for emerging epidemiological risks in the UK, *Culex pipiens* has been implicated as the primary vector of USUV transmission in England (Lawson et al., 2022). The detection of *Culex* hybrid forms thus poses a potential risk for disease spillover to humans. While the impact on human health is understood to be low (Cadar & Simonin, 2023), the first fatal case in an immunocompromised patient was recently recorded in Hungary (Szabó et al., 2025), underscoring the potential health risks of USUV in specific circumstances. Hybrids were detected sporadically from East to West in wetland, agricultural and urban settings across the entire study period, though *Cx. p. molestus* was not detected at any point during the study. This exemplifies a potential risk for human-biting *Culex* biotypes to act as bridge vectors to humans. Importantly, *Cx. p. pipiens* may still play a role in disease spillover to humans given its plasticity in feeding behaviour (Blom et al., 2024; Wehmeyer et al., 2024). These results corroborate evidence from northern European populations of *Cx. pipiens* s.l., which display clear genetic differentiation between above-ground and below-ground populations with relatively few hybrids (Haba & McBride, 2022). Therefore, an increased focus towards below-ground habitats may be needed to uncover ‘pure’ *molestus* biotypes in Northern Ireland. The differences in detections of *Cx. torrentium* between survey types (larvae vs adult) is also distinctive, but matches previous experimental work in other regions demonstrating the bias of adult baited traps towards *Cx. pipiens* and lack of efficacy at successfully attracting *Cx. torrentium* (Hesson et al., 2015). Nevertheless, we caveat that every adult sample of *Culex* was identified to species/biotype by PCR, whereas only a subset of larval samples of *Culex* were tested, meaning that *Cx. torrentium* could have been more frequent in larval collections than reported here. In our study, *Cx. torrentium* appeared to be more closely associated with urban land uses, being detected most frequently as larvae in artificial containers, with the few detections as adults tending to be in urban/agricultural settings. *Culex torrentium* were similarly only detected in artificial containers in the Republic of Ireland (Schaffner et al., 2024), albeit based only on three sampling events. Notably, a recent nationwide survey in England and Wales indicated that *Cx. torrentium* may have a preference to rural habitats over urban habitats at both adult and larval stages (Widlake et al., 2025), and in the wider European context, prefers to overwinter in ‘natural’ hibernacula (Sauer et al., 2023). The finding that Irish populations of *Cx. torrentium* may have preferences for oviposition in anthropogenic habitats is therefore novel, and requires further research.

Our study has several limitations which should be addressed in future studies. Firstly, surveillance was carried out over a single year, across a relatively short seasonal window (May to August). Previous work in England has demonstrated that the active period for *Culex pipiens* s.l. mosquitoes may span from April to October (Ewing et al., 2019), so future work should aim to address the full period of seasonal activity. Additionally, there may be interannual differences in mosquito ecology driven by temperature or rainfall variation which could only be captured via long-term surveillance over multiple years (*e.g.,* Simonetto et al., 2026). Also, our study relied on only one adult mosquito sampling method (BG-Pro traps), with a recent study in Scotland indicating that other traps types (*e.g*., Mosquito Magnet) are more sensitive to other mosquito species (*e.g*., *Anopheles;* Lee et al., 2026). Lastly, as this study focused on mosquito diversity and distribution, to understand the epidemiological risks from mosquito presence and abundance, future studies should aim to assess vector-host contact patterns and the competence of local vector populations to transmit disease.

## Conclusion

This study is the first systematic survey of mosquito vector ecology in wetlands and urban settings on the island of Ireland. We have demonstrated the widespread abundance and distribution of potential zoonotic vectors throughout Northern Ireland, particularly of *Culex pipiens pipiens*, the major vector of WNV and USUV in Europe. Further, we have provided evidence of potential hybridisation between the two *Culex pipiens* biotypes: a potential precursor to disease spillover to humans due to potential indiscriminate feeding by hybrid forms on birds and humans. Continued surveillance may be necessary to confirm the presence of the *Culex pipiens molestus* biotype, particularly towards below-ground habitats. Thus, we have exemplified a major risk factor to the emergence of mosquito-borne disease in Northern Ireland, namely through the widespread occurrence of potential vector species and suitability of wetland habitat. We further uncovered two previously unrecorded species for the island: *Culiseta fumipennis* and *Anopheles messeae*. Future work should continue to address geographical gaps around mosquitoes and associated pathogens in Northern Ireland, particularly as environments change, and enhance partnerships in One Health disease surveillance across the island of Ireland.

## Ethical approval

Ethical approval to sample in residential gardens sought and approved by Faculty of Medicine, Health and Life Sciences Research Ethics Committee (Faculty REC) in accordance with the Proportionate Review process (MHLS 25_81). Approval to conduct work on BANN ESTUARY AREA OF SPECIAL SCIENTIFIC INTEREST (ASSI) and LARNE LOUGH AREA OF SPECIAL SCIENTIFIC INTEREST (ASSI) was sought and approved by DAERA.

## Author contributions

R.M.C. – Conceptualisation, Data curation, Formal analysis, Investigation, Methodology, Visualisation, Writing – original draft

G.K. – Conceptualisation, Formal analysis, Investigation, Methodology, Visualisation, Writing – review & editing

L.S. – Investigation, Writing – review & editing

A.D. – Investigation, Writing – review & editing

J-P.P. – Supervision, Writing – review & editing

F.B. – Supervision, Writing – review & editing

E.P. – Conceptualisation, Funding acquisition, Methodology, Supervision, Writing – review & editing

J.T.A.D. – Conceptualisation, Funding acquisition, Methodology, Supervision, Writing – review & editing

H.M.F. – Conceptualisation, Funding acquisition, Methodology, Supervision, Writing – original draft

R.N.C. – Conceptualisation, Funding acquisition, Methodology, Supervision, Writing – original draft

## Acknowledgments

We thank the Wildfowl and Wetlands Trust, National Trust, Translink, Agri-Food and Biosciences Institute, Armagh City, Banbridge & Craigavon Council, Northern Ireland Water, Northern Ireland Environment Agency, Belfast Zoo and all other land managers and members of the public for permission to conduct fieldwork on their sites. We thank Meshach Lee for sharing equipment and advice for conducting fieldwork, David Kerrigan and Iona Latto for support in the laboratory, and Lucy Devlin and Christopher Mellon for support in the field. We thank all members of the Mosquito Scotland and Mosquito NI teams for their support.

