## Supplementary Information for "First nationwide mosquito surveillance in Northern Irish wetlands reveals presence of *Culex pipiens*/*molestus* hybrids"

**Table S1:** Details of field sites, target habitat(s) and sampling effort, organised by core sites and by additional snapshot sites. Where additional adult traps were deployed at a single site, this is indicated. All surveillance undertaken in private gardens has been anonymised to postcode level. Sampling effort was spread across May (n = 17), June (n = 18), July (n = 31) and August (n = 32). Adult collections were made near ponds (n = 20), reedbeds (n = 17), saltmarshes (n = 15), urban settings close to artificial containers (n = 16), wet grasslands (n = 15) and wet woodlands (n = 15), where n is the total number of adult traps deployed per month/target habitat type. Enhanced surveillance (*i.e.*, deployment of additional traps) was undertaken in July and August 2025 at the following sites: Ballycarry, Castle Archdale Country Park, Castle Archdale ICW, Oxford Island Nature Reserve.

| **Site name** | **County** | **Coordinates** | **Habitat(s)** | **Months visited (number of visits)** | **Number of adult traps** | **Number of aquatic habitats sampled** | **Nearest weather station (miles from site)** |
| --- | --- | --- | --- | --- | --- | --- | --- |
| **Core sites** | | | | | | | |
| Altarichard | Antrim | 55.10124, -6.26530 | Wet woodland | May – August (4) | 1-2 | 1 | Portglenone (24) |
| Ballycarry | Antrim | 54.77694, -5.72495 | Saltmarsh, wet woodland | June – August (3) | 2 | 2 | Killylane (15) |
| Ballykelly ICW | Derry | 55.04986, -7.02245 | Reedbed | May – August (4) | 1 | 1 | Magilligan #2 (7.8) |
| Ballykelly Village | Derry | 55.06251, -6.92088 | Urban garden | June – August (3) | 1 | 1 | Magilligan #2 (8.1) |
| Bangor City | Down | 54.65725, -5.68191 | Urban garden | May – August (5) | 1-2 | 2 | Helen’s Bay (3.3) |
| Beltrim | Tyrone | 54.72200, -7.23942 | Pond | May – August (4) | 1 | 1 | Castlederg (16) |
| Brookeborough | Fermanagh | 54.33667, -7.37264 | Wet woodland | May – August (4) | 1 | 1 | Thomastown (10) |
| Castle Archdale Country Park | Fermanagh | 54.47294, -7.73376 | Wet grassland, pond | May – August (4) | 1-2 | 2 | Thomastown (14) |
| Castle Archdale ICW | Fermanagh | 54.47399, -7.72060 | Reedbed | May – August (4) | 1-3 | 1 | Thomastown (14) |
| Castle Espie WWT | Down | 54.53077, -5.69544 | Pond | May – August (4) | 1 | 1 | Stormont Castle (8.3) |
| Clogher | Tyrone | 54.41920, -7.17823 | Wet grassland | May – August (4) | 1 | 1 | Thomastown (21) |
| Craigavon | Armagh | 54.47830, -6.36580 | Pond | May – August (4) | 1 | 1 | Armagh (18) |
| Gortin | Tyrone | 54.65632, -7.14010 | Urban garden | June – August (3) | 1 | 1 | Castlederg (19) |
| Loughgall | Armagh | 54.40607, -6.5995 | Wet woodland | May – August (4) | 1 | 1 | Armagh (7.9) |
| Oxford Island Nature Reserve | Armagh | 54.16470, -6.37165 | Wet grassland | May – August (4) | 1-2 | 1 | Armagh (19) |
| Portstewart | Derry | 55.16470, -6.75437 | Saltmarsh, artificial container | June – August (3) | 1 | 2 | Magilligan #2 (17) |
| Stoneyford ICW | Antrim | 54.56735, -6.11232 | Reedbed | May – August (4) | 1-2 | 1 | Stormont Castle (17) |
| Strangford Lough | Down | 54.53292, -5.69905 | Saltmarsh | May – August (4) | 1 | 1 | Stormont Castle (8.3) |
| **Snapshot sites** | | | | | | | |
| Balloo Wetland | Down | 54.63688, -5.66909 | Reedbed | May (1) | 0 | 1 | Helen’s Bay (4.3) |
| Ballydugan | Down | 54.31467, -5.74232 | Wet grassland, artificial container | July – August (2) | 2 | 3 | Katesbridge (18) |
| Belfast Zoo | Antrim | 54.65470, -5.94354 | Artificial container | August (1) | 1 | 1 | Stormont Castle (11) |
| Bog Meadows | Down | 54.58453, -5.96586 | Pond | August (1) | 0 | 1 | Stormont Castle (8.1) |
| Bratwell | Derry | 55.11676, -6.80214 | Pond | August (1) | 1 | 1 | Magilligan #2 (7.4) |
| Clabby | Fermanagh | 54.39727, -7.37241 | Reedbed | May (1) | 1 | 1 | Thomastown (12) |
| Donaghadee | Down | 54.63744, -5.56841 | Urban garden, pond | August (1) | 1 | 2 | Helen’s Bay (9.4) |
| Hillsborough | Down | 54.45208, -6.07395 | Wet woodland | July (1) | 1 | 0 | Katesbridge (17) |
| Larchfield | Down | 54.46242, -5.97326 | Pond | July – August (2) | 1 | 2 | Stormont Castle (16) |
| Moira | Down | 54.48467, -6.22839 | Pond | July (1) | 1 | 0 | Katesbridge (16) |
| Murlough | Down | 54.24647, -5.82973 | Saltmarsh, artificial container | July (1) | 2 | 2 | Murlough (0.1) |
| Stranmillis | Down | 54.56818, -5.94654 | Urban garden | July – August (2) | 1 | 0 | Stormont Castle (6.7) |

**Table S2:** Primer sequences used in this study.

| **Name** | **Primer sequence (5’ to 3’)** | **Reference** |
| --- | --- | --- |
| pipCQ11R | CATGTTGAGCTTCGGTGAA | (Bahnck & Fonseca, 2006) |
| molCQ11R | CCCTCCAGTAAGGTATCAAC |  |
| CQ11F2 | GATCCTAGCAAGCGAGAAC |  |
| ACEpipF | GGAAACAACGACGTATGTACT | (Smith & Fonseca, 2004) |
| ACEtorrF | TGCCTGTGCTACCAGTGATGTT |  |
| B1246sR | TGGAGCCTCCTCTTCACGG |  |
| 5.8S-UNF | TGTGAACTGCAGGACACATG | (Proft et al., 1999) |
| AATR | CGTTTGGCTTGGGTTATGA |  |
| AMSR | GACGCCTCACGATGACCTT |  |
| ADAR | CTTTCAATGCAAGAAATGGAT | (Kronefeld et al., 2014) |

**Table S3:** Summary of microclimatic conditions from April to August 2025 per weather station and associated field sites.

| **Weather station** | **Associated field sites** | **Mean temperature (°C) ± standard deviation** | **Total rainfall (mm)** |
| --- | --- | --- | --- |
| Armagh | Craigavon, Loughgall, Oxford Island Nature Reserve | 14.6 ± 3.29 | 265 |
| Castlederg | Beltrim, Gortin | 13.8 ± 3.42 | 425 |
| Helen’s Bay | Balloo Wetland, Bangor City, Donaghadee | 14.4 ± 3.04 | 292 |
| Katesbridge | Ballydugan, Hillsborough, Moira | 13.1 ± 3.69 | 333 |
| Killylane | Ballycarry | 12.9 ± 3.27 | 342 |
| Magilligan #2 | Ballykelly ICW, Ballykelly Village, Bratwell, Portstewart | 13.9 ± 3.48 | 346 |
| Murlough | Murlough | 14.2 ± 3.25 | 344 |
| Portglenone | Altarichard | 14.0 ± 3.31 | 356 |
| Stormont Castle | Belfast Zoo, Bog Meadows, Castle Espie WWT, Larchfield, Stoneyford ICW, Strangford Lough, Stranmillis | 14.4 ± 3.10 | 232 |
| Thomastown | Brookeborough, Castle Archdale Country Park, Castle Archdale ICW, Clabby, Clogher | 14.0 ± 3.13 | 387 |

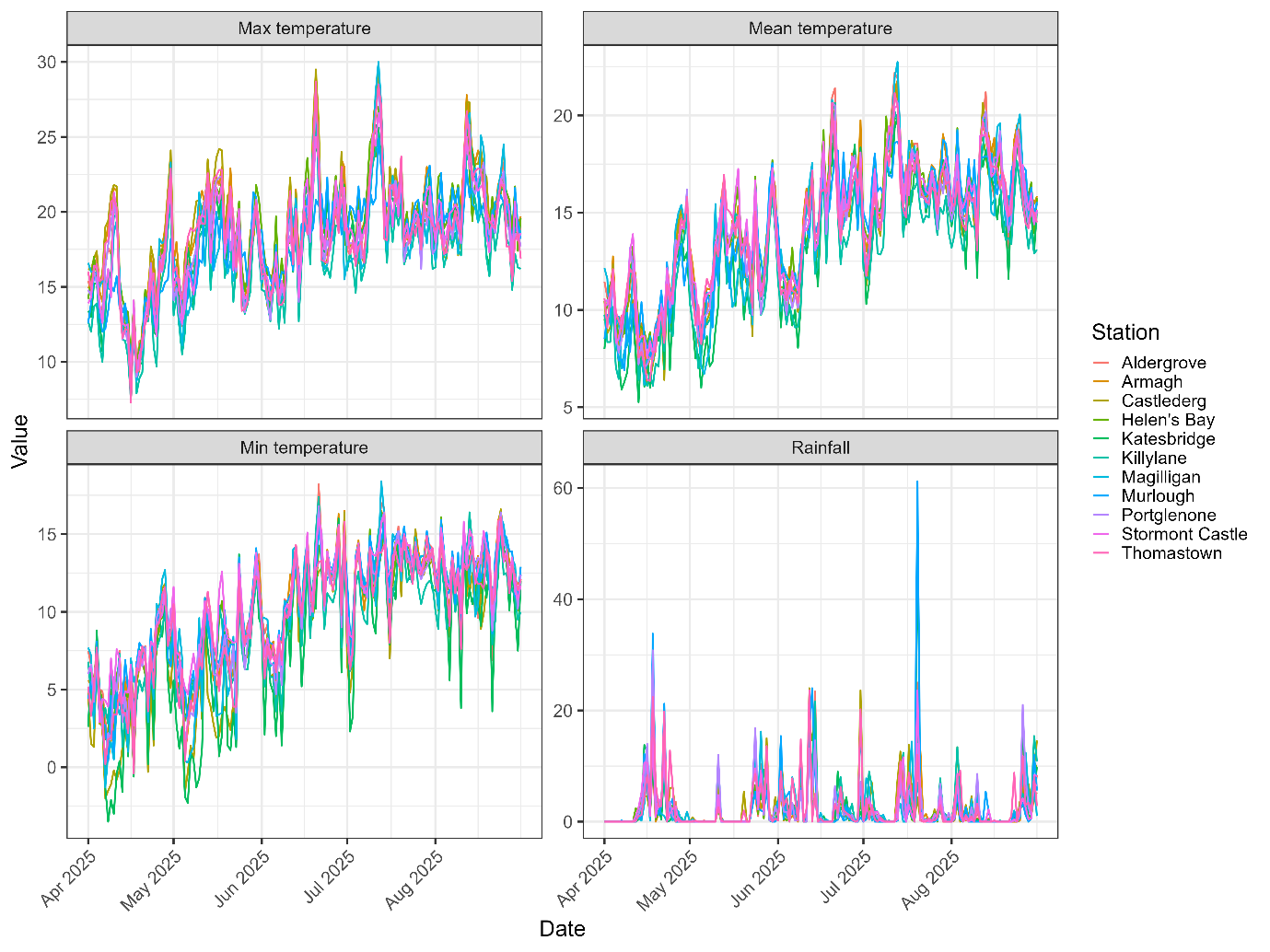

**Figure S1:** Summary of daily microclimatic conditions recorded at nearby weather stations to sampled wetlands in Northern Ireland during the study period (1^st^ April – 31^st^ August 2025).

**Table S4:** Mean temperature by month comparing the 1991-2020 30-year average Met Office and compiled weather station 2025 data.

| **Month** | **1991-2020 Met Office mean temperature (°C) ± standard deviation** | **2025 weather stations mean temperature (°C) ± standard deviation** |
| --- | --- | --- |
| April | 7.93 ± 1.09 | 10.1 ± 2.36 |
| May | 10.51 ± 0.93 | 12.4 ± 2.40 |
| June | 13.02 ± 0.95 | 14.6 ± 2.81 |
| July | 14.66 ± 0.99 | 16.4 ± 2.02 |
| August | 14.48 ± 0.91 | 16.3 ± 1.74 |

**Table S5:** Sampling and ecological data by species sampled as larvae across the 41 collections where mosquito larvae were detected, in descending order based on abundance. Here, *Culex pipiens* s.l. includes *Culex pipiens pipiens*, *Culex pipiens molestus*, *Culex pipiens/molestus* hybrids and *Culex torrentium*, as not every larval specimen of *Culex* was tested by PCR to species/biotype level. Total number of collected specimens (N), total number of collections where species was present (P), temperature range and mean (T °C), pH range and mean (pH) and associated habitats with number of collections per habitat type (n).

| **Species** | **N** | **P** | **T °C (mean)** | **pH (mean)** | **Associated habitats (n)** |
| --- | --- | --- | --- | --- | --- |
| *Culex pipiens* s.l. | 1139 | 25 | 15.4-24.5 (18.4) | 6.65-10.6 (7.70) | Reedbed (5), pond (3), artificial container (10), ditch (3), saltmarsh (1), wet woodland (1), wet grassland (2) |
| *Aedes detritus* (Haliday) | 279 | 9 | 17.8-26.7 (21.78) | 6.71-7.23 (7.04) | Saltmarsh (9) |
| *Culiseta annulata* (Schrank) | 47 | 8 | 12.2-17.9 (16.04) | 6.65-7.84 (7.34) | Reedbed (3), pond (2), wet woodland (3) |
| *Culiseta morsitans* (Theobald) | 9 | 1 | 15.4 | 7.03 | Ditch (1) |
| *Anopheles claviger* (Meigen) | 3 | 2 | 16.3-17.4 (16.85) | 6.95-8.49 (7.72) | Pond (2) |
| *Anopheles maculipennis* s.l. (Meigen) | 2 | 2 | 18.0-21.7 (19.85) | 6.88-7.75 (7.32) | Pond (2) |

**Table S6:** Summary of statistical significance of variables tested in this study, resulting from generalised linear mixed models. The single *Culex* spp. larval mosquito presence model was analysed using a binomial distribution, whereas all abundance models were analysed using negative binomial distributions. Significant results are in **bold**.

| **Variable** | **χ^2^** | **Df** | **p-value** |
| --- | --- | --- | --- |
| **Abundance of larval mosquitoes** |  |  |  |
| Water temperature | 3.327 | 1 | 0.068 |
| Ph | 0.870 | 1 | 0.351 |
| Habitat depth | 0.134 | 1 | 0.714 |
| Shade | 0.125 | 2 | 0.939 |
| Habitat size | 0.439 | 2 | 0.803 |
| Vegetation coverage | 21.046 | 3 | **<0.001** |
| Predator presence | 8.754 | 1 | **<0.01** |
| Month | 0.160 | 3 | 0.984 |
| Wetland type | 26.930 | 6 | **<0.001** |
| **Presence of *Culex* spp. larval mosquitoes** |  |  |  |
| Water temperature | 17.625 | 1 | **<0.01** |
| Habitat depth | 0.052 | 1 | 0.819 |
| Habitat size | 0.415 | 2 | 0.813 |
| Predator presence | 11.345 | 1 | **<0.001** |
| Wetland type | 5.841 | 6 | **0.016** |
| **Abundance of *Culex pipiens/torrentium* larval mosquitoes** |  |  |  |
| Water temperature | 0.456 | 1 | 0.499 |
| Habitat depth | 0.151 | 1 | 0.698 |
| Habitat size | 7.292 | 2 | **0.026** |
| Predator presence | 7.269 | 1 | **<0.01** |
| Month | 9.837 | 3 | **0.020** |
| **Abundance of adult mosquitoes** |  |  |  |
| Mean temperature | 0.088 | 1 | 0.767 |
| Total rainfall during trapping | 0.116 | 1 | 0.733 |
| Lagged rainfall (1-week) | 3.976 | 1 | **0.046** |
| Lagged rainfall (2-weeks) | 0.210 | 1 | 0.647 |
| Lagged rainfall (3-weeks) | 0.987 | 1 | 0.320 |
| Lagged rainfall (4-weeks) | 1.224 | 1 | 0.269 |
| Latitude | 1.640 | 1 | 0.200 |
| Longitude | 1.829 | 1 | 0.176 |
| Month | 19.847 | 3 | **<0.001** |
| Wetland type | 26.117 | 5 | **<0.0001** |
| **Abundance of *Culex pipiens/torrentium* adult mosquitoes** |  |  |  |
| Mean temperature | 3.578 | 1 | 0.060 |
| Total rainfall during trapping | 2.220 | 1 | 0.136 |
| Lagged rainfall (1-week) | 4.221 | 1 | **0.040** |
| Lagged rainfall (2-weeks) | 0.304 | 1 | 0.581 |
| Lagged rainfall (3-weeks) | 0.354 | 1 | 0.552 |
| Lagged rainfall (4-weeks) | 3.843 | 1 | **0.050** |
| Latitude | 5.867 | 1 | **0.015** |
| Longitude | 3.179 | 1 | 0.075 |
| Month | 15.236 | 3 | **<0.01** |
| Wetland type | 25.748 | 5 | **<0.0001** |
| **Abundance of *Culiseta annulata* adult mosquitoes** |  |  |  |
| Mean temperature | 0.015 | 1 | 0.902 |
| Total rainfall during trapping | 0.100 | 1 | 0.760 |
| Lagged rainfall (1-week) | 3.459 | 1 | 0.063 |
| Lagged rainfall (2-weeks) | 0.763 | 1 | 0.382 |
| Lagged rainfall (3-weeks) | 1.183 | 1 | 0.277 |
| Lagged rainfall (4-weeks) | 2.604 | 1 | 0.107 |
| Latitude | 0.362 | 1 | 0.548 |
| Longitude | 0.650 | 1 | 0.420 |
| Month | 2.667 | 3 | 0.446 |
| Wetland type | 22.03 | 5 | **<0.001** |

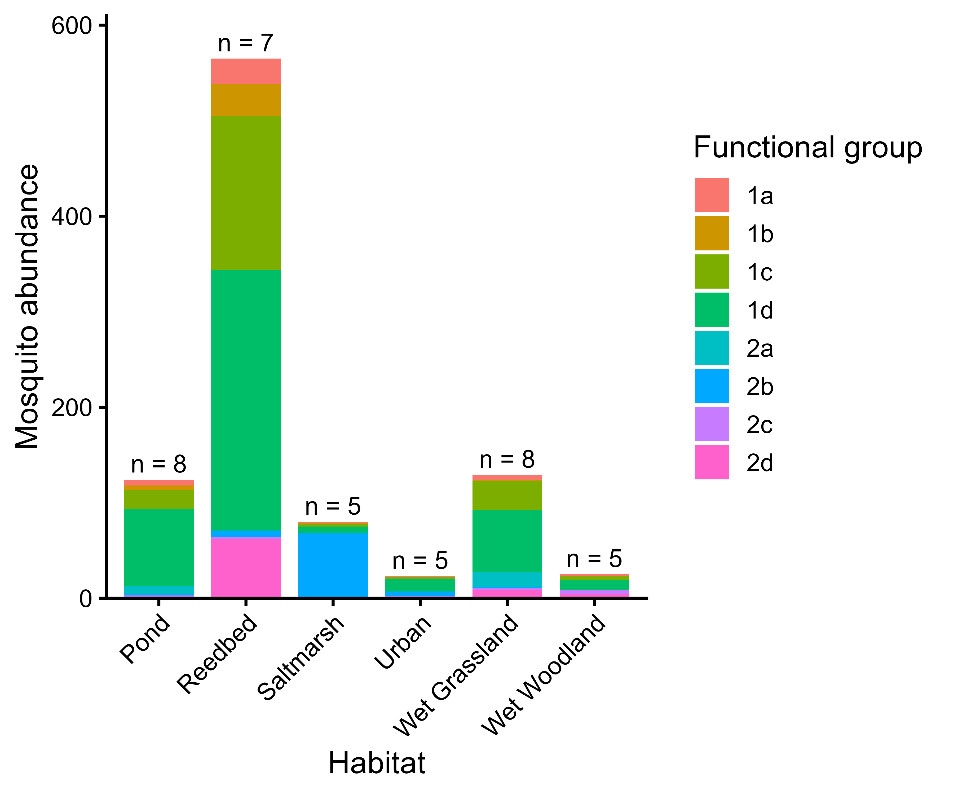

**Figure S2:** Total abundance of mosquitoes by functional group per wetland habitat type as defined by Medlock et al. (2024). The number of functional groups present in the given wetland habitat type is indicated above each bar.

**Text S1: Notes on the use of squishing buffer to extract DNA from larval *Culex* mosquitoes**

The success rate for successfully amplifying target DNA from collected field samples of *Culex* differed between adult (96% success) and larval (53% success) surveys. It was suspected that the reason for the lower amplification success rate of larval *Culex* was due to the differences in sample preservation. Specifically, if larval *Culex* were not processed in a timely manner and preserved in ethanol prior to mortality, this could reduce the efficacy of the DNA extraction method.

To test this, larval mosquito samples (n=20) were collected from laboratory colonies of *Culex pipiens pipiens* (Belfast) and *Culex torrentium* (Bangor). The amplification success rate was 100% for both species when larvae were directly killed in ethanol, while the success rate was 60% and 20% for *Culex pipiens pipiens* and *Culex torrentium* respectively when DNA was extracted from larvae already found to be dead in the rearing pan.

Thus, it may be concluded that the lower success rates for the amplification of larval *Culex* DNA may be due to inadequate preservation methods that are likely to be incompatible with our DNA extraction technique.

**Text S2: Description of natural history of mosquito species previously collected in Ireland but not in the present study**

In our study, we detected 15 of the previously recorded 21 Irish mosquito taxa. During our study, we did not record presence of either adult or larval stage *Aedes punctor, Aedes cinereus, Aedes geminus, Aedes dorsalis, Anopheles algeriensis* and *Anopheles daciae*. Previous detections of *Aedes punctor, Aedes cinereus* and *Aedes geminus* have largely been limited to the South of Ireland in County Cork (*Aedes punctor*, Ashe et al. (1991)) and County Kerry (all three species, Ashe et al. (1991); Schaffner et al. (2024)). *Ae. dorsalis* is a coastal-breeding species which has not been recorded on the island of Ireland since its detection in 1929 in County Wicklow (Ashe et al., 1991). We hypothesise that these four species (*Aedes cinereus, Aedes geminus, Aedes punctor* and *Aedes dorsalis*) may potentially be limited to a southern distribution on the island, given the lack of historical and contemporary records of these species in Northern Ireland, though expanded surveillance over further sites and seasons will be key to determining this. *Anopheles algeriensis* has previously been recorded in Central Ireland in County Clare (Ashe et al., 1991) and County Offaly (Schaffner et al., 2024), though was not collected in the present study. *Anopheles daciae* has only recently been confirmed on the island through molecular analyses on one occasion in County Galway (Schaffner et al., 2024), though the *An. maculipennis* complex has been documented widely throughout the island (Ashe et al., 1991). Continued surveillance may confirm the presence of *Anopheles daciae* in Northern Ireland, and of *Anopheles atroparvus* on the island of Ireland.
